# Protein language models and the long tail of functional diversity

**DOI:** 10.64898/2026.08.14.744703

**Authors:** Ria Vinod, Samir Char, Ava P. Amini, Lorin Crawford, Kevin K. Yang

**Affiliations:** Center for Computational Molecular Biology, Brown University, Providence, RI, USA; Microsoft Cloud & AI, Redmond, WA; Microsoft Research, Cambridge, MA, USA

## Abstract

Protein language model performance on downstream tasks depends on the pretraining data, motivating recent efforts to combine genomic- and metagenomic-derived protein sequences into large-scale atlases. Because these datasets are highly redundant, sequences are typically clustered by similarity and sampled during training. Sequences that do not belong to any cluster, known as “singletons”, are typically excluded from training and evaluation because they are considered to be artifacts. However, singletons represent the long tail of functional diversity and are abundant in many large-scale atlases: nearly 43% of the 3.34 billion sequences in the joint genomic-metagenomic dataset GigaRef are singletons. Here, we characterize singletons derived from UniRef and GigaRef by assessing whether clustering missed homologs, how much their exclusion affects protein language model (PLM) training, and which biological domains they contain. We find that many GigaRef singletons belong to a cluster under alternative parameter settings, suggesting that genomic and metagenomic datasets may require dataset-specific clustering configurations. We also show that singletons share mutual information with clustered sequences, making them learnable by PLMs and useful for training. Finally, metagenomic singletons carry denser, more diverse domain content than clustered sequences, including domain-level homology that sequence-identity clustering misses. Together, these results support including singletons in PLM training and call for closer examination of data curation in large-scale integrated sequence atlases.

## Introduction

Protein language models (PLMs) are powerful tools for learning patterns within the vast biological diversity captured in sequence datasets, and they have become widely used for structure prediction [1], variant effect prediction [2], protein engineering [3–6], and sequence design [7]. Because standard masked or autoregressive PLMs are trained to minimize cross-entropy when reconstructing amino-acid sequences, their performance on downstream tasks is fundamentally dictated by the composition of their training data [8, 9]. This sensitivity to training dataset composition has motivated the field to expand datasets to cover wider evolutionary and sequence diversity [10–13], under the assumption that broader coverage can improve model generalization and translate to improved performance on downstream biological tasks.

To increase the sequence diversity of model training datasets, recent efforts have combined metagenome-derived proteins with existing protein sequence collections spanning organisms from diverse clinical and environmental settings [10–13]. These large-scale datasets integrate genomic resources such as UniRef [14] with metagenomic databases [15–22], thereby encompassing proteins derived from both genomes and metagenomes. To maximize diversity and minimize redundancy, the standard approach is to cluster sequences on the basis of sequence identity, grouping those above chosen identity and coverage thresholds into clusters that are then sampled from during training [11, 23]. Proteins that have insufficient sequence identity to any other cluster form single-member clusters, i.e., “singletons”.

Metagenomic singletons are typically excluded from model training and evaluation. For example, 43% of the 3.34 billion sequences in GigaRef, a combined genomic and metagenomic atlas, are singletons. This practice reflects the assumptions that these proteins may arise from sequencing noise or assembly errors and that models may struggle to learn sequences lacking evolutionary neighbors in the training set [8, 13, 24]. However, singletons may capture the long tail of functional diversity, as novel proteins that are unlikely to cluster with other proteins are also ones that are likely to contain new domains, be sampled from new evolutionary lineages, and have new functions. In addition, recent work has demonstrated that singletons are partly an artifact of imprecise sequence-level clustering, as a subset can be rescued into existing clusters via similarity in a PLM’s representation space [10].

In this work, we analyze singletons in genomic (UniRef [25]; Fig. 1A) and metagenomic (GigaRef [12]; Fig. 1B) protein sequence datasets and investigate their behavior during PLM training (Fig. 1C-E). First, an exhaustive search recovered singletons missed by the original clustering and revealed substantially different recovery rates between the genomic and metagenomic regimes, indicating distinct clustering failure modes (Fig. 1F). We then trained a PLM on GigaRef-singletons and performed a large-scale, sequence-level inference study of PLM training dynamics on singletons and their clustered counterparts (Fig. 1G). Excluding singletons from training hurts PLM performance, and singletons share mutual information with clustered sequences that PLMs can learn from during training. Finally, we correlated sequence-level PLM learnability with Pfam [26] domain content to determine which biological domains PLMs learn across training data compositions. Singletons contained a broader range of domains than clustered sequences, and PLMs successfully learned from a substantial subset of them (Fig. 1H). Taken together, our analyses indicate that singletons should not be discarded as a uniform category, but characterized, partially recovered, and selectively included in both PLM training and evaluation pipelines.

**Figure 1.**
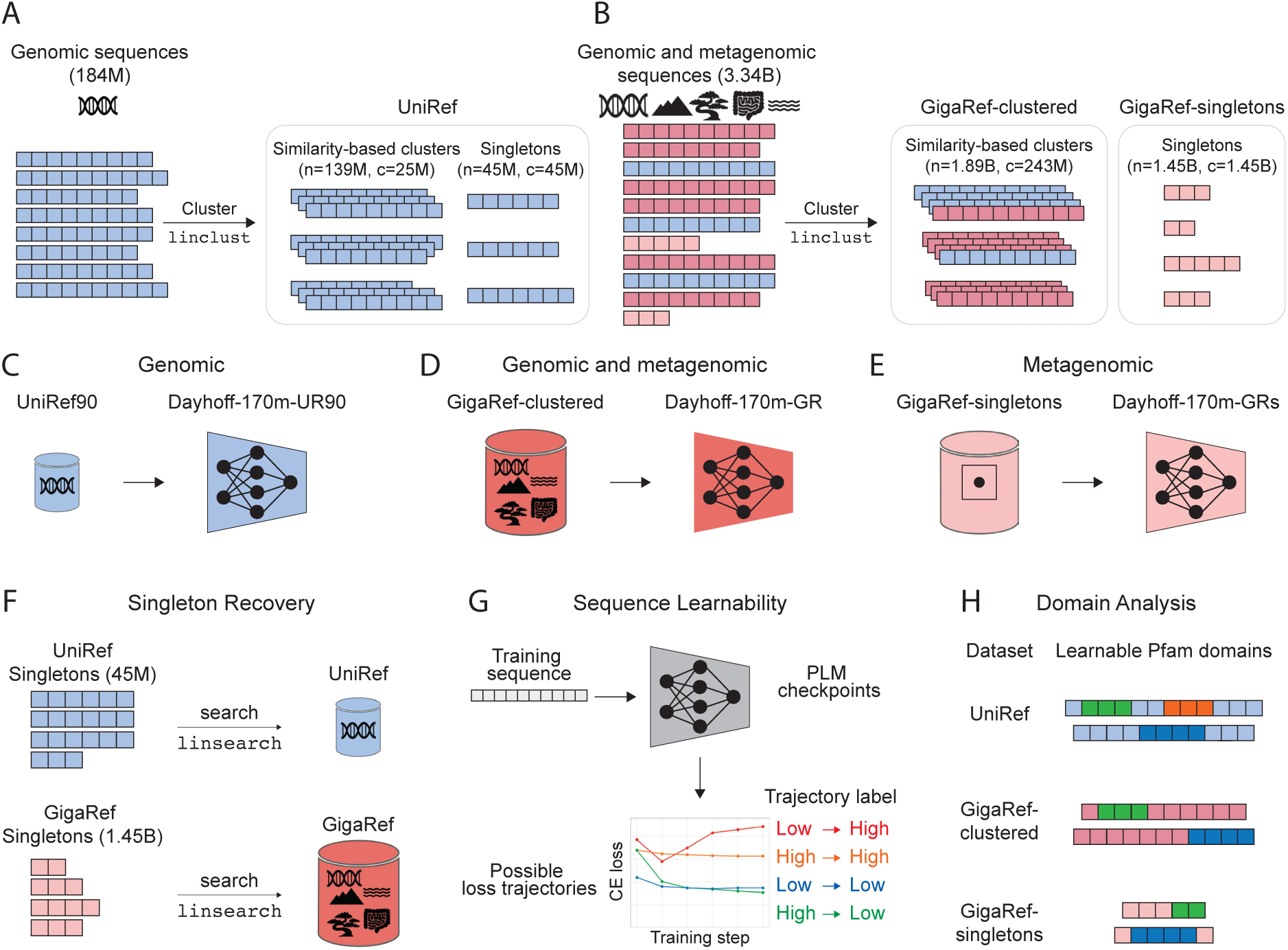
Overview of genomic and metagenomic dataset construction, trained protein language models, and subsequent analyses. Clustering a protein sequence dataset creates multi-member and single-member clusters (i.e., singletons). **(A)** UniRef clustering algorithm. **(B)** GigaRef clustering algorithm. **(C-E)** Three Dayhoff-170m protein language models are trained on datasets spanning the genomic-to-metagenomic regime: UniRef90 (genomic-derived), GigaRef-clustered (genomic- and metagenomic-derived), and GigaRef-singletons (metagenomic-derived), yielding PLMs (C) Dayhoff-170m-UR90, (D) Dayhoff-170m-GR, and (E) Dayhoff-170m-GRs, respectively. **(F)** Nearest-neighbor search of UniRef- and GigaRef-singletons against their respective full databases can identify homologs missed by the original clustering protocol. **(G)** Training sequences can be classified as *learnable* or *not learned* by a PLM based on their loss trajectories over training. **(H)** Pfam domain annotation of sequences in each dataset can be compared with sequence learnability to identify learnable and not learned domains.

## Results

### Many singletons are not really singletons

We first asked whether the sequences designated as singletons are true singletons, or whether some have homologs in the dataset that the original clustering failed to identify. Both UniRef and GigaRef were clustered with MMseqs2 linclust [27]. UniRef clusters were generated by first clustering UniRef100 at 90% sequence identity, and the resulting UniRef90 cluster representatives were then re-clustered at 50% identity. This final 50% clustering step produced 19 million UniRef50 clusters (spanning 139 million UniRef90 sequences) and 45.1 million singletons (Fig. 1A). Throughout, we refer to these two sets as UniRef-clustered and UniRef-singletons, respectively. GigaRef was constructed using an analogous hierarchical clustering procedure, but with additional pre-clustering to accommodate its larger scale. Specifically, sequences from nine genomic and metagenomic resources were combined [14, 16–22]; because the total input exceeded the sequence limit of MMseqs2, MERC and SRC were first combined and pre-clustered at 70% identity, while MGnify was independently pre-clustered at 70% identity. These representative sequences were then combined with sequences from SMAG, MetaEuk, MGV, GPD, TOPAZ, and UniRef100 and clustered at 90% identity, and the resulting representatives were re-clustered at 50% identity to produce GigaRef (Fig. 1B). Throughout, we refer to the 1.89 billion sequences in multi-member clusters and the 1.45 billion singletons at the 50% threshold as GigaRef-clustered and GigaRef-singletons, respectively. Under these protocols, UniRef consists of 24.4% singletons and GigaRef consists of 43.4% singletons.

Across the four groups, mean sequence length decreases steadily from UniRef-clustered (367 amino acids) to UniRef-singletons (264 amino acid) to GigaRef-clustered (192 amino acid) to GigaRef-singletons (149 amino acid), with spread narrowing in the same order: GigaRef-singletons are both the shortest and the most uniform in length (Fig. 2A). MMseqs2 assigns a sequence to a cluster only if its alignment to the cluster representative satisfies both a minimum sequence-identity threshold and a coverage threshold; sequences that do not satisfy both against any representative are retained as singletons. In --cov-mode 1, the coverage is computed with respect to the query’s length, in --cov-mode 2, the coverage is computed with respect to the length of the cluster representative, which is, by definition, the longest sequence in the cluster. Both UniRef and GigaRef were clustered under coverage mode 2 at 80% coverage and required the alignment to span at least 80% of the cluster representative’s length (Fig. 2B). Because the representative is the longest sequence in each candidate group, this is a length-asymmetric criterion: a short candidate cannot satisfy the 80% requirement against a substantially longer representative, regardless of how well their overlapping regions align or if they satisfy the criteria against a shorter sequence within the cluster. This suggests that a single clustering criterion may not be appropriate for genomic and metagenomic datasets, which have vastly different sequence characteristics.

**Figure 2.**
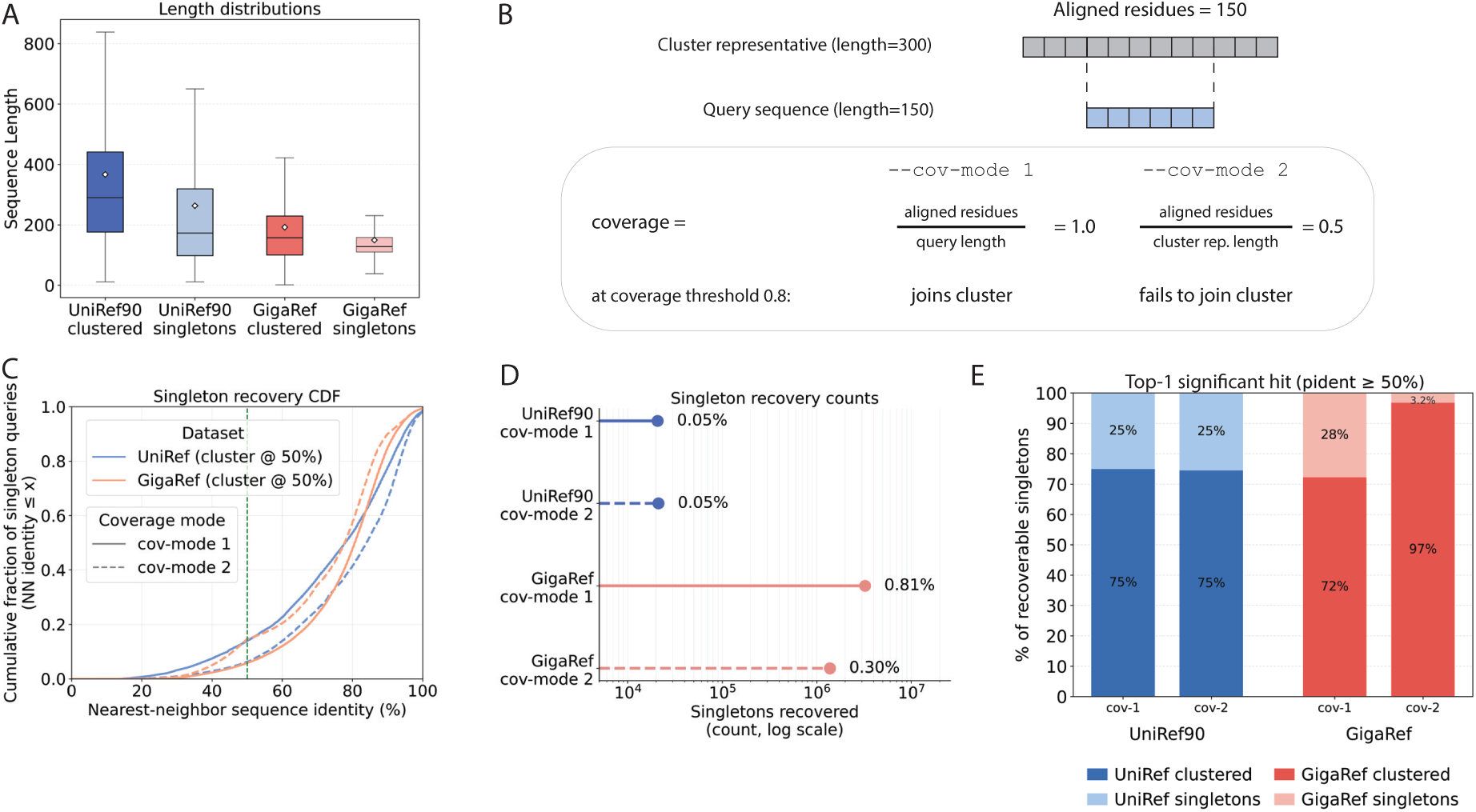
Recovery of false singletons via search. **(A)** Sequence length distributions for singleton and clustered sequences in UniRef (blue hues) and GigaRef (pink hues). Boxes span the interquartile range (IQR) with the median shown as a horizontal line and the mean as an open diamond; whiskers extend to 1.5*×*IQR. **(B**) Overview of MMseqs2 cluster assignment at 80% coverage fraction: coverage mode 1 normalizes aligned residues by query length, coverage mode 2 by cluster representative length. **(C**) Cumulative fraction of singleton queries (y-axis) versus top-1 significant hit sequence identity in percentage (x-axis) for UniRef-singletons (blue) and GigaRef-singletons (pink). **(D)** Percentage of UniRef90 and GigaRef singleton queries with a significant hit in the corresponding dataset with at least 50% sequence identity. **(E)** Fraction of recoverable singletons that have significant hits in clusters vs. singletons.

To test this directly, we queried each dataset’s singletons against all of the sequences in the original dataset (prior to clustering) using linsearch [27]. Note that linsearch is more thorough than linclust: it searches each query against the full database rather than only against cluster current representatives. By keeping the coverage mode fixed to that of the original linclust, we can measure the effect of varying the search criterion independently of the rest of the pipeline. We ran linsearch on all 45 million UniRef-singletons and on a subset of 400 million GigaRef-singletons against their respective full databases under both coverage mode 2 at 80% coverage (matching the original clustering criterion) and also coverage mode 1 at 80% coverage (which requires the alignment to span 80% of the shorter sequence in the linsearch pair rather than the representative). Because linsearch compares only sequence pairs that share at least one k-mer (the same prefilter linclust uses), any false-singleton rate we report is a lower bound: singletons whose true homologs share no k-mers with them are not detected by this recovery procedure.

We consider a singleton to be “recoverable” if it has at least one significant linsearch hit in the full dataset above the original 50% clustering identity threshold. Across recoverable singletons, the most significant hit was often a close relative, with the cumulative distributions for both UniRef and GigaRef lying overwhelmingly above the 50% cutoff (Fig. 2C). A singleton passing this cutoff is false by linclust’s own criteria: it satisfies the identity and coverage thresholds against a cluster member, but linclust only tests sequences against cluster representatives, so the qualifying alignment was never evaluated.

Despite low recovery rates, the absolute number of false singletons is large: under coverage mode 2 at 80% coverage, linsearch recovered homologs for 1,249,467 GigaRef-singletons (0.26% of the 400 million subset queried; Table S1) and 21,260 UniRef-singletons (0.04% of 45,148,521; Table S2). Switching to coverage mode 1 at the same coverage threshold approximately tripled GigaRef to 3,271,392 recovered singletons (0.76%), while leaving UniRef90 essentially unchanged (20,817 still equating to about 0.04%) (Fig. 2D). This coverage-mode asymmetry tracks the length composition of each dataset. UniRef90, composed of longer genomic proteins with more uniform lengths, is largely insensitive to the sequence length used to compute coverage. In contrast, GigaRef’s shorter, more heterogeneous metagenomic proteins are recovered far more effectively under cov-mode 1, where a short query needs to cover only itself rather than a longer representative sequence. Among the recovered singletons, the majority matched an existing multi-member cluster rather than another singleton, confirming that they are misclassified members of known clusters rather than true singletons (Fig. 2E).

Extrapolated across all 1.45 billion GigaRef-singletons, the 0.76% rate corresponds to *∼* 1.1 *×* 10^7^ length-orphaned false singletons—each a homolog of a clustered sequence but retained as a singleton likely because it was too short to span 80% of a longer representative. At this scale, even sub-1% recovery produces millions of false singletons. Furthermore, this establishes that a substantial fraction of singletons in metagenomic-integrated atlases are not true biological orphans but artifacts of clustering criteria that behave differently across the genomic and metagenomic regimes.

Finally, we asked whether singletons share information with sequences in multi-member clusters using a compression-based test. The mutual information between two data streams appears as the savings gained by compressing them together rather than separately [28]. We drew 50 matched pairs of subsamples, one of singletons and one of cluster representatives (*∼* 2.6 *×* 10^7^ residues each), and compressed each pair separately versus jointly (see Methods). For both UniRef90 and GigaRef, joint compression of singletons and clustered sequences was smaller than the sum of compressing each subset apart, indicating that singletons share reusable sequence structure with clustered proteins (Fig. S1).

### Singletons are important for PLM training

We first examined whether singletons are less learnable than other sequences. A training sequence’s loss under a PLM is correlated with the size of cluster to which it belongs in both the genomic (Fig. 3A) and metagenomic (Fig. 3B) regimes; singletons have the highest losses. However, using the final loss alone obscures how productive the sequence was as a training data point. We thus did a deeper learnability analysis that examined how individual sequence losses evolved over the course of training for Dayhoff-170m-UR90, Dayhoff-170m-GR, and Dayhoff-170m-GRs. Following the framework introduced in Lin et al. [29], we consider a training input learnable if its loss meaningfully decreases over the course of training (High*→*Low). Conversely, a sequence is not learned if its loss does not meaningfully reduce over training–either remaining high (High*→*High) or even increasing (Low*→*High). An input may also be trivial for the model to learn if the loss is low at the onset of training and remains low throughout (Low*→*Low).

**Figure 3.**
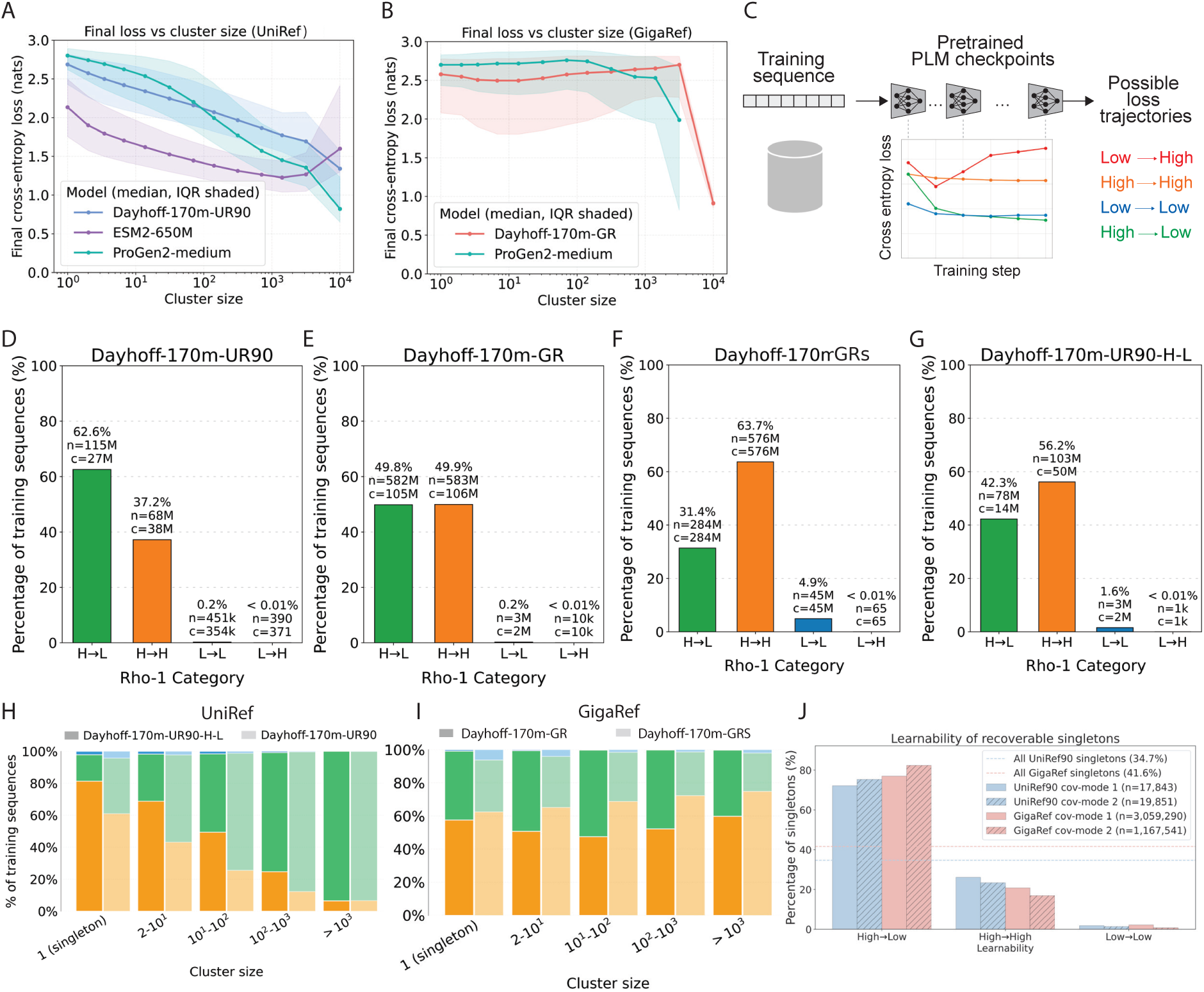
Analysis of sequence learnability for PLMs trained on UniRef90, GigaRef, and GigaRef-singletons. **(A)** Cluster size (x-axis) vs. final sequence loss (y-axis) for UniRef90 training sequences under Dayhoff-170m-UR90 (blue), ESM-2 650M (purple), and ProGen2-medium (teal). **(B)** Cluster size (x-axis) vs. final GigaRef-training sequence loss (y-axis) for GigaRef sequences under Dayhoff-170m-GR (orange) and ProGen2-medium (teal). **(C)** A sequence learnability framework based on loss trajectories. **(D–G)** Fraction of training data by loss trajectory label for (D) Dayhoff-170m-UR90, (E) Dayhoff-170m-GR, (F) Dayhoff-170m-GRs, and (G) Dayhoff-170m-UR90-H-L. On each panel, *c* denotes the number of clusters spanned by *n* sequences in each category of trajectories. **(H)** Comparison of UniRef90 sequence learnability under Dayhoff-170m-UR90 and Dayhoff-170m-UR90-H-L, binned by cluster size. **(I)** Comparison of GigaRef sequence learnability under Dayhoff-170m-GR and Dayhoff-170m-GRs, binned by cluster size. **(J)** Learnability of recoverable singletons vs. all singletons. Grouped bars give the percentage of recoverable query singletons falling into each learnability category.

For each training sequence, we evaluated its loss at six checkpoints, from the untrained PLM through the fully trained model, and classified the resulting trajectory into one of the four learnability categories (Fig. 3C). We then fit a least-squares line to the six log-loss values across the checkpoints and summarized its trajectory by the net change Δ*L* = *L*_end_ *− L*_start_. Each sequence was then sorted into one of the four learnability categories (High*→*Low, High*→*High, Low*→*Low, Low*→*High) by comparing Δ*L* to a threshold *τ* . The threshold *τ* separates trajectories that meaningfully change from those that remain stagnant: sequences with Δ*L < −τ* decreased in loss and were labeled High*→*Low, while those with Δ*L > τ* increased and were labeled Low*→*High. For sequences whose losses do not meaningfully change (*|*Δ*L| ≤ τ* ), we used the final loss relative to the dataset mean to distinguish Low*→*Low (final loss below the mean) from High*→*High (final loss above the mean); see Methods for details. We analyzed the sequence-level training dynamics of three PLMs identical in architecture and training objective: Dayhoff-170m-UR90, trained on 184 million genomic-derived proteins in UniRef90 (sampled uniformly by UniRef50 cluster); Dayhoff-170m-GR, trained on 1.17 billion genomic- and metagenomic-derived proteins in GigaRef-clustered (sampled uniformly by cluster) [12]; and Dayhoff-170m-GRs, trained only on 905 million metagenomic-derived GigaRef-singletons.

We first analyzed the training sequences of Dayhoff-170m-UR90 (see example learnable and not learned sequences in Table S3). Although the final training sequence perplexity was 10.3 (indicative of a successfully trained PLM), only 62.6% of sequences were learnable (Fig. 3D; Table S4). The training sequence perplexity of Dayhoff-170m-GR was 9.8, lower than that of Dayhoff-170m-UR90. However, only 49.8% of training sequences in GigaRef-clustered were learnable (Fig. 3E; Table S5). Dayhoff-170m-GRs had a training sequence perplexity of 12.3, and only 31.4% of training sequences were learnable (and notably, 4.9% Low*→*Low), while 63.7% were not learned (Fig. 3F; Table S6). There was more variation across GigaRef sequence loss trajectories than across UniRef90 trajectories, consistent with more diversity in the training data (Fig. S2A-B) and a similarly larger skew in High*→*High cluster loss means (Fig. S3A-B). Additionally, even though Dayhoff-170m-GR was not trained on GigaRef-singletons, the model was able to successfully reduce loss on many singleton sequences, with a mean sequence perplexity of 10.7 on GigaRef-singletons (Table S7), providing further evidence that GigaRef-singletons share mutual information with clustered sequences. Likewise, Dayhoff-170m-GRs achieved a mean sequence perplexity of 12.4 on GigaRef-clustered sequences (Table S8). Although PLMs tend to achieve lower loss on longer sequences, Low*→*Low and High*→*High classified sequences have similar length distributions, suggesting that learning is not only driven by length (Fig. S4).

While the learnability analyses revealed that some fraction of singletons are learnable, they also revealed that some clustered sequences are not learned. We then asked whether excluding sequences that were not learned from training would improve model learning, a direct comparison to prior work [29] that showed that not all tokens are equally useful during pretraining. We trained Dayhoff-170m-UR90-H-L on only the UniRef90 sequences that Dayhoff-170m-UR90 classified as High*→*Low, which removed 68 million High*→*High sequences, including 45.1 million singletons, from training. We then re-evaluated learnability under this new model. Surprisingly, we found that restricting training to previously learnable sequences did not keep them learnable: the training sequence perplexity of 12.3 was worse than that of Dayhoff-170m-UR90 (10.3), and only 42.3% of these originally High*→*Low sequences remained High*→*Low under Dayhoff-170m-UR90-H-L (Fig. 3G). When we compared per-sequence learnability under the two models, binned by the size of the cluster to which each sequence belongs, we observed that removing the High*→*High sequences lowered learnability across many cluster sizes and not just the singletons (Fig. 3H; Table S9). We repeat this comparison between Dayhoff-170m-GR and Dayhoff-170m-GRs, contrasting the effect of holding out all GigaRef-singletons against training on only GigaRef-singletons (Fig. 3I; Table S10).

Finally, we asked whether the recoverable singletons–those for which linsearch identified a significant nearest neighbor–are also the sequences that PLMs can learn. We analyzed the learnability of recoverable singletons across both UniRef and GigaRef in both coverage modes (Fig. 3J). Learnability among recovered singletons (72.1% vs. 75.0% for UniRef90 and 77.0% vs. 82.4% for GigaRef, under coverage modes 1 vs. 2, respectively) far exceeded that of the full singleton pool prior to recovery filtering (34.7% for UniRef90 and 41.6% for GigaRef). These results were consistent regardless of the dataset they were derived from and which clustering protocol designated them as singletons, showing that singletons with a remote homolog in the training data are consistently learnable.

### Singletons have detectable domain homology

Sequence-level loss trajectories tell us whether a sequence is learned but not what biology that sequence encodes. Because protein domains are the subunits that make up sequences and dictate the biology they encode, we sought to characterize the biological domains underlying singleton and clustered populations and their degree of learnability by a PLM. We used Pfam [26] to annotate the domains present in each sequence and also the domains present in most significant hits of the recoverable singletons (Fig. 4A). Among 121,000 randomly selected GigaRef-singletons, the rate of detectable Pfam domain homology was largely independent of nearest-neighbor sequence identity: even singletons whose most significant hit falls below the 50% clustering threshold contained detectable domains (Fig. 4B). This persistence of homology across the clustering boundary is consistent with the limited sensitivity of linear clustering itself: DIAMOND DeepClust reports that MMseqs2/linclust groups only 21.6% of sequences sharing a Pfam domain architecture, as opposed to 68.6% for sensitive all-versus-all clustering [30]. The link between domain homology and learnability is reflected in the singletons a PLM learns (High*→*Low), which carry detectable Pfam hits more often (61%) than those it fails to learn (High*→*High, 37%) or the small Low*→*Low set (under 2%) (Fig. 4C). No singletons in this analysis were classified as Low*→*High.

**Figure 4.**
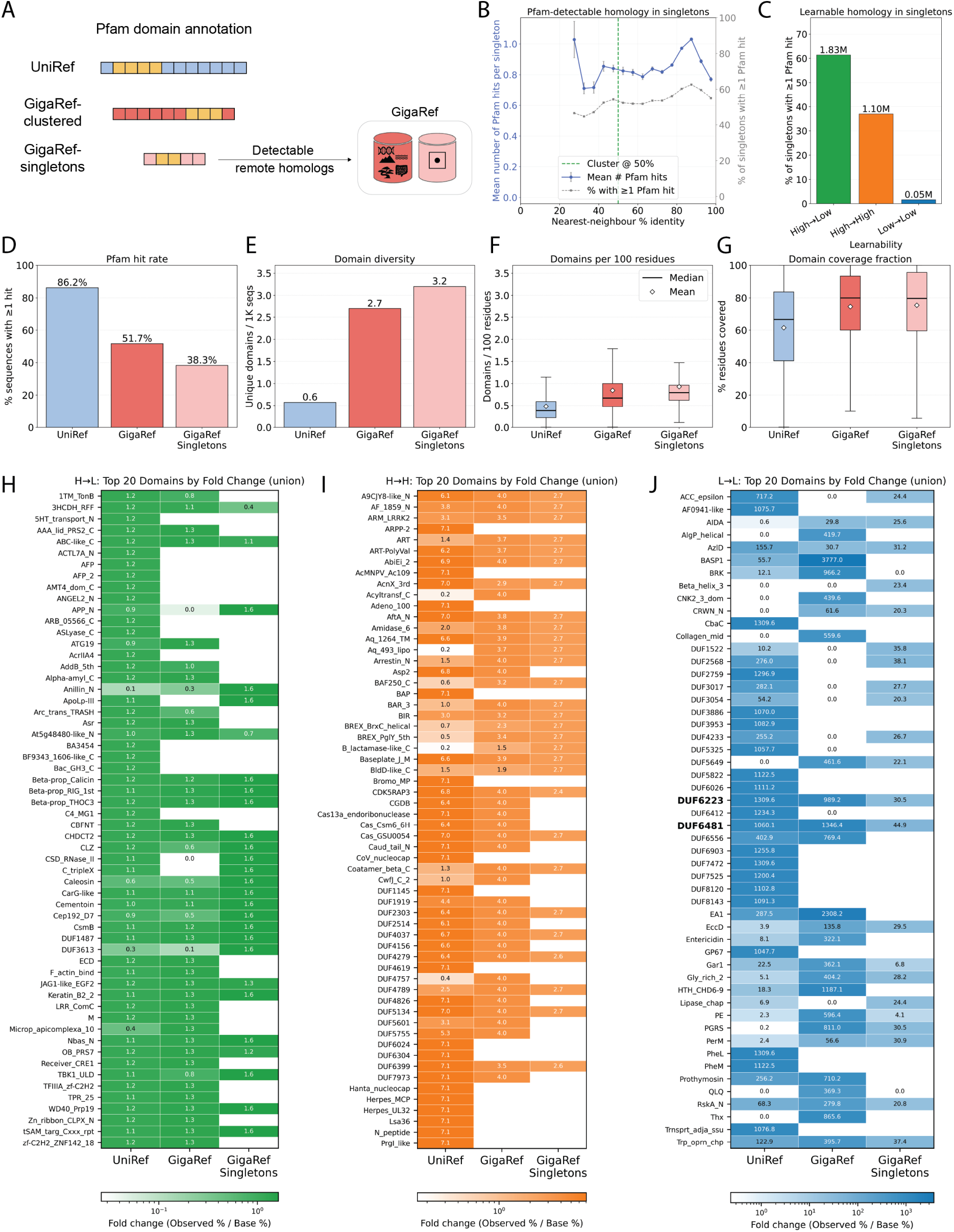
Pfam domain and enrichment analysis. **(A)** Pfam hidden Markov models (HMMs) annotate biological domains across sequences. **(B)** Pfam-detectable homology in singletons as a function of nearest-neighbour sequence identity. **(C)** Pfam domains across loss trajectory categories (High*→*Low, High*→*High, Low*→*Low). **(D)** Pfam hit rate (percentage of sequences with *≥*1 domain hit) across UniRef90, GigaRef-clustered, and GigaRef-singleton datasets. **(E)** Domain diversity (unique domains per 1,000 sequences) across datasets. **(F)** Domains per 100 residues across datasets. Box plots show interquartile range (IQR); whiskers extend to 1.5*×*; lines and diamonds denote median and mean, respectively. **(G)** Domain coverage fraction across datasets; box plots as in (F). **(H–J)** Cross-dataset overlap of the most strongly enriched Pfam domains by loss-trajectory category. For each category and each of the three datasets, Pfam domains were ranked by fold change (observed % */* base %), and the top 20 significantly enriched domains (Benjamini–Hochberg corrected Fisher exact *p <* 0.05 and fold change *>* 1) were retained; each heatmap shows the union of these domains across the three datasets. Cell color intensity (log scale) and cell text both encode the fold change for that category–dataset pair. Bold domain names are those appearing in the top 20 of all three datasets. Blank cells indicate the domain was not tested in that dataset (too few total hits to enter the enrichment analysis); white cells annotated “0.0” indicate the domain occurs in the dataset but in no sequence of that category.

If singletons were predominantly sequencing artifacts or pipeline-induced false positives, their annotated content should be sparser and less domain-rich than genomic and clustered metagenomic sequences. While the Pfam hit rate per sequence falls with dataset diversity (Fig. 4D): 86.2% of UniRef90 sequences had at least one Pfam hit, compared to 51.7% of GigaRef-clustered sequences and 38.3% of GigaRef-singleton sequences, this is entirely due to sequence length. The number of unique domains per 1,000 sequences rose from 0.6 in UniRef90 to 2.7 in GigaRef and 3.2 in Singletons (Fig. 4E), and the domain density per 100 residues was higher in GigaRef-clustered and GigaRef-singletons (median *∼*0.7 in both) than in UniRef90 (median *∼*0.5; Fig. 4F). The domain coverage fraction was similarly elevated in GigaRef and GigaRef-singletons relative to UniRef90 (Fig. 4G). Thus, the dataset-level Pfam profile demonstrates that singletons capture a distinct functional landscape landscape.

We next stratified Pfam annotations by learnability category and, within each category and each dataset, ranked domains by fold change–the ratio of the percentage of category sequences containing a domain to that domain’s dataset-wide base rate (observed % / base %)–and retained the top 20 significantly enriched domains for each category (Benjamini–Hochberg-corrected Fisher exact *P <* 0.05, fold change *>* 1). We visualized the union of these per-dataset top-20 sets (i.e., the union of sets for each of UniRef, GigaRef, and GigaRef-singletons, per learnability category; Fig. 4H–J). Because fold change is bounded above by a category’s inverse prevalence (*N*_total_*/N*_category_), the attainable enrichment is small for the dominant High*→*Low class and large for the rare Low*→*Low class. The Low*→*High category was excluded because it contained a negligible number of sequences across all three datasets. There is almost no overlap in enriched domains between the three learnability categories (Fig. 4H–J).

The most enriched High*→*Low domains were almost entirely disjoint across datasets: of 58 domains in the union, *none* appeared in the top 20 of all three datasets, and the dataset leaders shared no members (e.g. 3HCDH RFF and AAA lid PRS2 C in UR90, LRR ComC and Keratin B2 2 in GR, and Nbas N and WD40 Prp19 in GRs; Fig. 4H). Likewise, the most enriched High*→*High domains were completely disjoint across datasets, but include many viral and defense-associated domains (Adeno 100, CoV nucleocap, the baculovirus domain AcMNPV Ac109, and the CRISPR effector Cas13a endoribonuclease in UR90; the phage and CRISPR domains Caud tail N, Cas Csm6 6H, and Cas GSU0054 in GigaRef).

The most enriched Low*→*Low domains are domains of unknown function (DUFs) and other rare families found almost exclusively in Low*→*Low sequences.

## Discussion

We performed the first large-scale study of singleton protein sequences to characterize the long tail of functional diversity in genomic and metagenomic datasets. We argue against the current status quo of discarding singletons as a uniform category by showing that (1) many singletons are falsely designated as singletons by improperly configured clustering protocols, (2) singletons can be successfully learned by a PLM and are indeed important during training, and finally, (3) metagenomic singletons encode the most diverse domains per sequence, and their residues are more likely to fall within an annotated domain when compared to genomic sequences. We selectively recover and characterize a dataset of singletons for PLM training and evaluation, offering the opportunity to improve PLM performance on various downstream tasks by exposing the models to the long tail of functional sequence diversity. Our clustering and Pfam analyses together indicate that recoverable singletons are frequently sequence fragments–partial reads whose truncated length explains both their failure to meet coverage thresholds during clustering and their designation as singletons–yet these fragments retain dense, diverse domain content and are learnable by a PLM. An abundance of fragments is a characteristic of large-scale integrated sequence atlases [27, 31, 32]. As efforts to expand and categorize sequence diversity have grown, the computational and memory demands of clustering at scale have driven the adoption of increasingly heterogeneous protocols, and the cost of re-clustering makes these choices effectively permanent. Today, public and proprietary genomic and metagenomic datasets are clustered in widely varying ways–differing in their pre-clustering and filtering steps, cascading clustering schemes, coverage modes and thresholds, sequence-identity cutoffs, minimum cluster membership, and conventions for handling singletons (Table S11). To our knowledge, this is the first in-depth study of singletons and the clustering protocols by which they were designated as singletons. We restrict our conclusions to UniRef90 and GigaRef for two reasons: both were clustered by the same protocol, enabling us to do a controlled comparison across the genomic and metagenomic regimes, and both have publicly accessible singletons. We also limit our sequence learnability analysis to the Dayhoff family of models, whose training checkpoints are publicly available.

However, our work highlights two gaps in the field. First, there has been little effort to study the effects of clustering protocols, but choices around clustering parameters tangibly matter. We show that singleton recovery on the basis of sequence identity has very different outcomes under different clustering parameters for genomic versus metagenomic datasets. As a result, while multiple datasets integrate common sources such as MGnify [16] and JGI [15], they arrive at very different cluster counts, with 207M non-singleton clusters in OMG [13], 243M non-singleton (1.70B total) clusters in GigaRef [12], and 1.1 billion clusters in ESM-C training. Second, singleton handling is similarly inconsistent across prior work. Sometimes singletons are removed, as in GLM2 [13], ProGen2 [11], and ProtGPT3 [33]. However, in ESM-C [10], it is not explicitly reported that singletons are left out of training, and subsequently singletons were found to cluster in latent space. Newer clustering algorithms such as DIAMOND DeepClust [30] push the sensitivity–speed frontier of large-scale clustering, enabling tens of billions of sequences to be clustered at low identity thresholds with high sensitivity. This expands the singleton problem: deep clustering 19 billion sequences at 30% identity designates 68% of them as singletons. More complete dataset and clustering releases would enable rigorous singleton characterization to guide the selection of training data.

Finally, the downstream consequences of identifying training data inputs from the long tail of functional diversity remains to be tested in the lab. Previous work has shown that training larger models on more diverse data improved protein expression rates for generated designs [10, 12, 34]. We expect that including singletons during training will aid in this goal. In sum, our work motivates the re-examination of clustering protocols as the field progresses toward larger scale data integration and model training and lays the foundation for the recovery and inclusion of the long tail of protein sequence functional diversity.

## Code and data availability

All code to reproduce the analyses will be available under an open-source MIT license at https://github.com/microsoft/dayhoff. Inference results and analysis outputs will also be made available via the Zenodo link in the repository. All Dayhoff-170m model checkpoints and training datasets used in this study are available at https://huggingface.co/collections/microsoft/dayhoff-atlas.

## Acknowledgments

This research was conducted using a combination of computational resources and services provided by Microsoft Research and the Center for Computation and Visualization at Brown University. The authors thank Sarah Alamdari for helpful discussions and Neil Tenenholtz and Sean Whitzell for assistance on using Microsoft’s compute resources.

## Author Contributions

**Ria Vinod**: Conceptualization, Methodology, Software, Formal Analysis, Investigation, Visualization, Writing – Initial Draft, Writing – Reviewing & Editing. **Samir Char**: Data Curation. Writing – Reviewing & Editing. **Ava P. Amini**: Supervision, Writing – Reviewing & Editing. **Lorin Crawford**: Resources, Supervision, Writing – Reviewing & Editing. **Kevin K. Yang**: Conceptualization, Methodology, Investigation, Supervision, Resources, Writing – Reviewing & Editing.

## Methods

### Compression rate of singletons versus clustered sequences

We measure how much a fixed classical compressor saves when given two sequence sets jointly rather than separately. Following the framing of language modeling as compression in Delétang et al. [28], the compressed length of a sequence set *X* under a fixed lossless compressor is denoted *L*(*X*). The redundancy this compressor exposes between two sets is

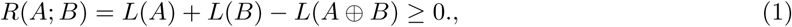

where *A ⊕ B* denotes the concatenation of *A* and *B*. In the limit of an optimal compressor, *L*(*·*) *→ H*(*·*) and *R*(*A*; *B*) *→ I*(*A*; *B*), where *H*(*·*) is the Shannon entropy of the source and *I*(*A*; *B*) denotes the mutual information between *A* and *B*. For any fixed practical compressor, *R* is a compressor-bounded surrogate of mutual information that captures only the redundancy the compressor can exploit. The fraction of bits saved by joint compression,

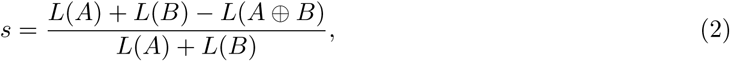

normalizes this redundancy by the separately-compressed total. Throughout, we use + between two sets to denote that they are compressed separately and the resulting bit lengths are summed, and ⊕ to denote that they are concatenated and compressed jointly in a single pass.

We instantiate this on two datasets. For each dataset *D ∈ {*UniRef90, GigaRef}, we let *A* be the set of singleton sequences (denoted UR90-S and GR-S) and *B* be one representative per non-singleton cluster (UR90-c and GR-c). Taking one representative per cluster prevents intra-cluster redundancy, which is high by construction, from dominating *L*(*B*) and trivially inflating the joint saving. We compress only the concatenated amino-acid residues (one byte per residue), discarding FASTA headers and newlines, so that measured redundancy reflects sequence content rather than shared metadata.

The compressor is xz -9e (LZMA preset 9 with the EXTREME flag), whose dictionary window is fixed at 64 MiB. We adopt a subsampling protocol to ensure that *A_i_*, *B_i_*, and *A_i_ ⊕ B_i_*, where *i* denotes the index of the replicate, each fit within a single window such that the joint compressor can in principle exploit cross-set structure. From each set we make a single streaming pass over the FASTA file, keeping each sequence with a probability from a Bernoulli distribution chosen so that the resulting candidate pool contains *∼* 4*S* residues, where *S* = 25 MB is the per-replicate residue budget. The factor of four balances pool-construction cost against the independence of subsequent draws. We then draw *N* = 50 replicates: for the *i*-th replicate, we sample sequences without replacement from the pool until *S* residues are accumulated, producing subsets *A_i_, B_i_* of sizes 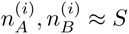. The pool itself is constructed once per set, so the across-replicate variance we report reflects within-pool subsampling variance rather than the full pipeline variance under re-drawing from the corpus.

Each of *A_i_*, *B_i_*, and *A_i_ ⊕ B_i_* is then compressed independently, yielding bit lengths 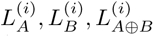 after subtracting the constant LZMA header (measured by compressing an empty stream). The order of sequences within *A_i_* and within *B_i_* is randomized per replicate, but *A_i_* always precedes *B_i_* in *A_i_ ⊕ B_i_*. Since LZMA is sequential, *L*(*A ⊕ B*) has a mild order dependence that this convention fixes consistently across replicates. Per replicate, we report the separate and joint compression rates

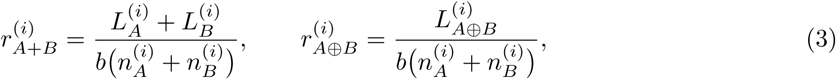

which are the compressed-bits-per-raw-bit when the two sets are processed separately and as a single concatenated stream, respectively. Here *b* = 8 bits/byte converts the raw residue byte counts 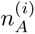 and 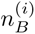 in the denominator to bits, matching the units of the bit-length numerators. Across the *N* = 50 replicates, we report the mean and standard deviation of 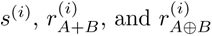.

### Sequence learnability: classifying sequences on the basis of training loss trajectory

Autoregressive PLMs are trained by applying a next-token prediction loss to all tokens in a protein sequence. Recent work has examined token-level training dynamics of language models, and shown that different tokens have different loss patterns, which can lead to training inefficiency and also hurt model performance [29]. In this work, we extend this framework to the sequence level and categorize sequences into four groups based on their loss trajectory over the course of model training: (1) High*→*High sequences with high perplexity at the start and end of training; (2) High*→*Low sequences with high initial perplexity that decreases over training; (3) Low*→*Low sequences with low perplexity throughout training; and (4) Low*→*High sequences whose perplexity increases over training. We interpret High*→*Low sequences as those the model successfully learns, Low*→*Low sequences as those the model finds trivially predictable from initialization, and High*→*High and Low*→*High sequences as those the model fails to learn (either by never reducing loss or by drifting toward higher loss as training progresses). These sequences are categorized by the following trajectory clustering protocol. Given a sequence of loss measurements (*ℓ*_0_*, ℓ*_1_*, . . . , ℓ_n_*) recorded at each of *n* + 1 training checkpoints, we fit a linear model by minimizing the sum of squared residuals

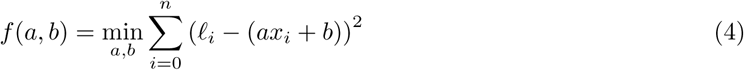

where *x_i_* corresponds to the training step at checkpoint *i*. Linear fitting provides robustness to noise at individual checkpoints and captures the overall trend for sequences with non-monotonic loss trajectories. From the fitted parameters, we compute the loss at the start and end of training

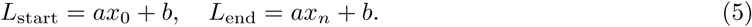

The change in loss over the training period is then:

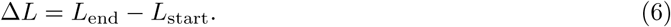

We additionally compute *L*_mean_, the mean final loss across all sequences in the dataset. Sequences are classified into four categories based on Δ*L* and the final loss *ℓ_n_*where

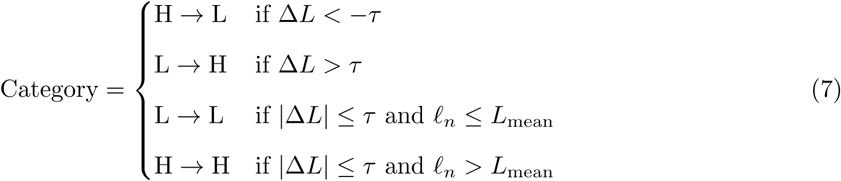

where *τ* is a threshold parameter controlling the sensitivity of the categorization. Sequences with *|*Δ*L| ≤ τ* are considered stagnant (i.e., their loss did not change meaningfully during training), and we distinguish between stagnant sequences that were always easy (Low*→*Low) versus always hard (High*→*High) by comparing their final loss *ℓ_n_* to the dataset mean *L*_mean_. For interpretability, final results are reported via the transformation perplexity = exp(*L*).

## Supplementary Tables

**Table S1.**
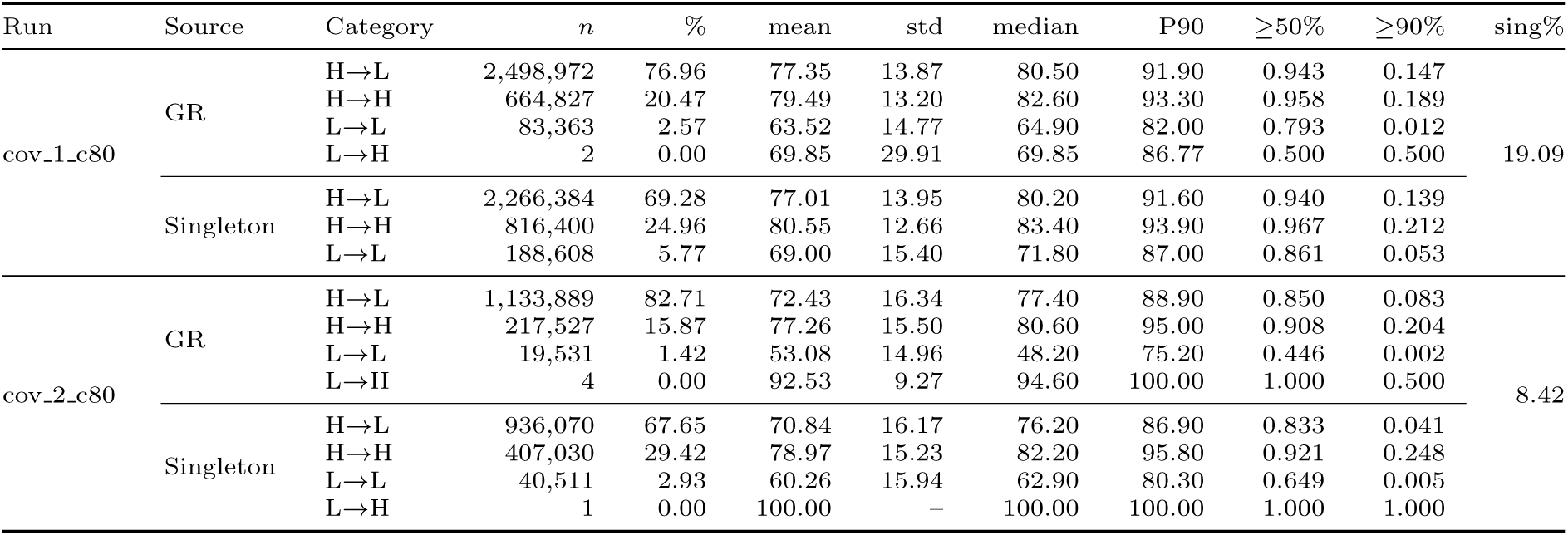
Nearest-neighbor statistics for GigaRef-singletons. Per-Rho-1-category breakdown of singleton queries with at least one linsearch nearest neighbor in GigaRef-Singletons (self-hits excluded). Learnability labels are assigned by the Dayhoff-170m-GR and Dayhoff-170m-GRs PLMs at *τ* = 0.2. Two coverage modes are reported at 80% coverage fraction: cov-mode 1 (400M singletons queried) and cov-mode 2 (443M queried). The % column gives each label’s share of queries with a significant nearest neighbor hit. Identity columns report the mean, standard deviation, median, and 90th percentile of pident across hits, plus the fraction of queries whose nearest neighbor has pident *≥* 50% and *≥* 90%. The final sing% column is the run-level fraction of NN target hits that are themselves GigaRef-singletons, i.e. singleton*/*(singleton+non-singleton).

**Table S2.** linsearch statistics for UniRef90 singletons. Per-learnability category breakdown of UniRef90 singleton queries with at least one linsearch hit in UniRef90-Singletons (self-hits excluded). Learnability labels are assigned by the Dayhoff-170m-UR90 PLM at *τ* = 0.2. Two coverage modes are reported at 80% coverage fraction: cov-mode 1 and cov-mode 2 (both 45M singletons queried). The % column gives each label’s share of queries with a significant hit. Identity columns report the mean, standard deviation, median, and 90th percentile of pident across hits, plus the fraction of queries whose most significant hit has pident *≥* 50% and *≥* 90%. The final sing% column is the run-level fraction of most significant target hits that are themselves UniRef90 singletons, i.e. singleton*/*(singleton+non-singleton). The L*→*H category is omitted as no sequences in it had a significant hit.

| Run | Category | *n* | % | mean | std | median | P90 | $\geq 50\%$ | $\geq 90\%$ | sing% |
| --- | --- | --- | --- | --- | --- | --- | --- | --- | --- | --- |
| cov_1.c80 | H→L | 15,522 | 74.89 | 71.01 | 19.93 | 75.00 | 93.70 | 0.829 | 0.185 | 15.81 |
|  | H→H | 4,829 | 23.30 | 81.69 | 14.20 | 85.50 | 96.80 | 0.964 | 0.355 |  |
|  | L→L | 375 | 1.81 | 71.17 | 19.10 | 73.80 | 94.00 | 0.845 | 0.211 |  |
| cov_2.c80 | H→L | 16,096 | 76.02 | 77.94 | 16.76 | 82.30 | 95.40 | 0.929 | 0.314 | 19.89 |
|  | H→H | 4,776 | 22.56 | 84.54 | 13.30 | 88.40 | 97.70 | 0.972 | 0.441 |  |
|  | L→L | 302 | 1.43 | 73.84 | 18.69 | 77.10 | 94.30 | 0.871 | 0.262 |  |

**Table S3.**
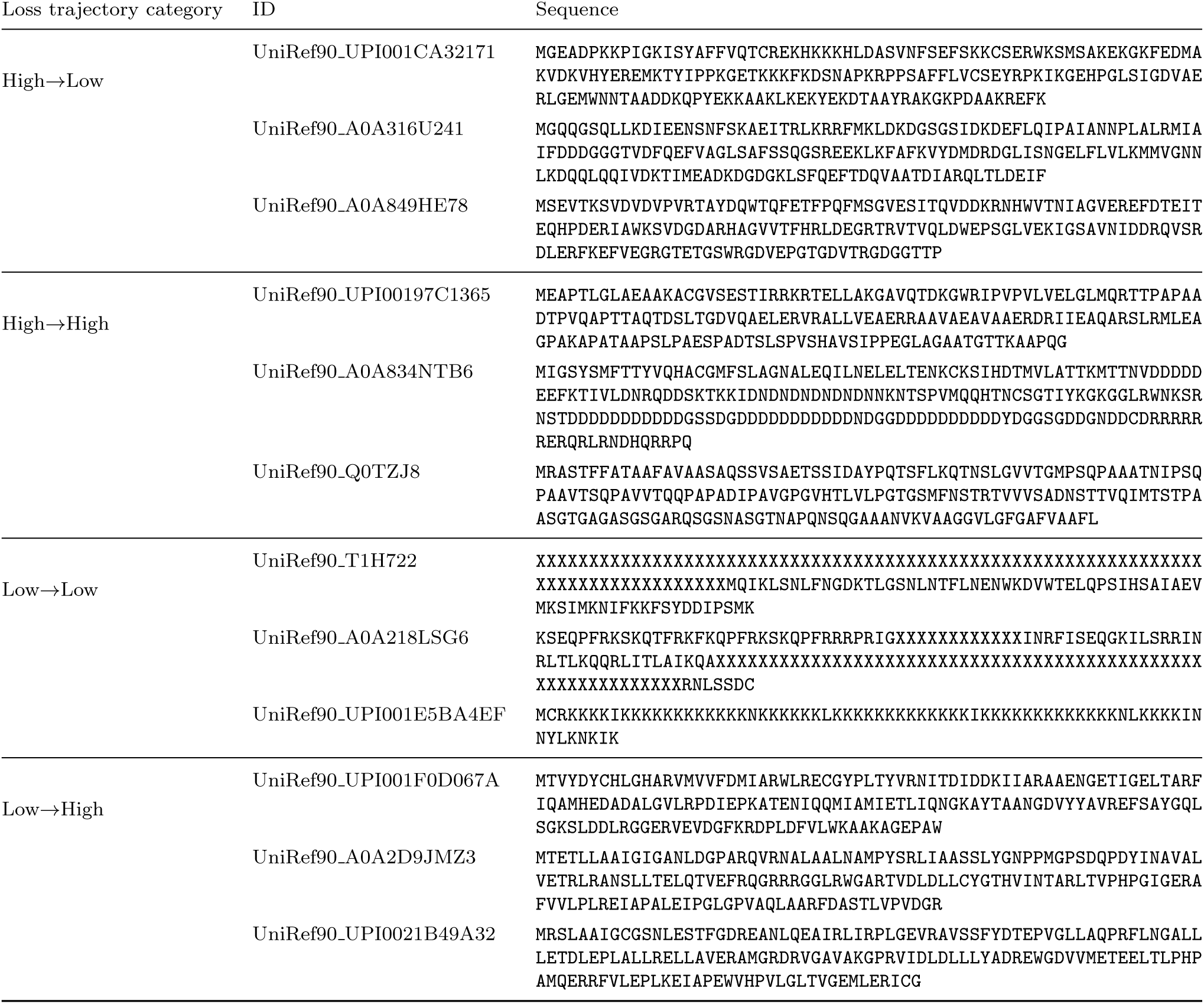
Representative sequences from the sequence learnability analysis for Dayhoff-170m-UR90.

**Table S4.** Learnability categorization of Dayhoff-170M-UR90 training sequences. Sensitivity analysis of threshold parameter *τ* on sequence categorization. Linear fit across 6 checkpoints {1000, 16000, 31000, 46000, 61000, 76000}; reference step 76,000. Global *L*_mean_ = 2.3340 (PPL 10.32). Total sequences: 183,900,910 across 207 shards.

| $\tau$ | Category | $N$ | % |
| --- | --- | --- | --- |
| 0.15 | H→L | 122,555,266 | 66.64 |
|  | H→H | 61,096,569 | 33.22 |
|  | L→L | 247,846 | 0.13 |
|  | L→H | 1,229 | 0.00 |
| 0.20 | H→L | 115,055,320 | 62.56 |
|  | H→H | 68,394,692 | 37.19 |
|  | L→L | 450,508 | 0.24 |
|  | L→H | 390 | 0.00 |
| 0.25 | H→L | 107,973,707 | 58.71 |
|  | H→H | 75,159,082 | 40.87 |
|  | L→L | 767,945 | 0.42 |
|  | L→H | 176 | 0.00 |
| 0.30 | H→L | 100,911,835 | 54.87 |
|  | H→H | 81,643,757 | 44.40 |
|  | L→L | 1,345,200 | 0.73 |
|  | L→H | 118 | 0.00 |

**Table S5.**
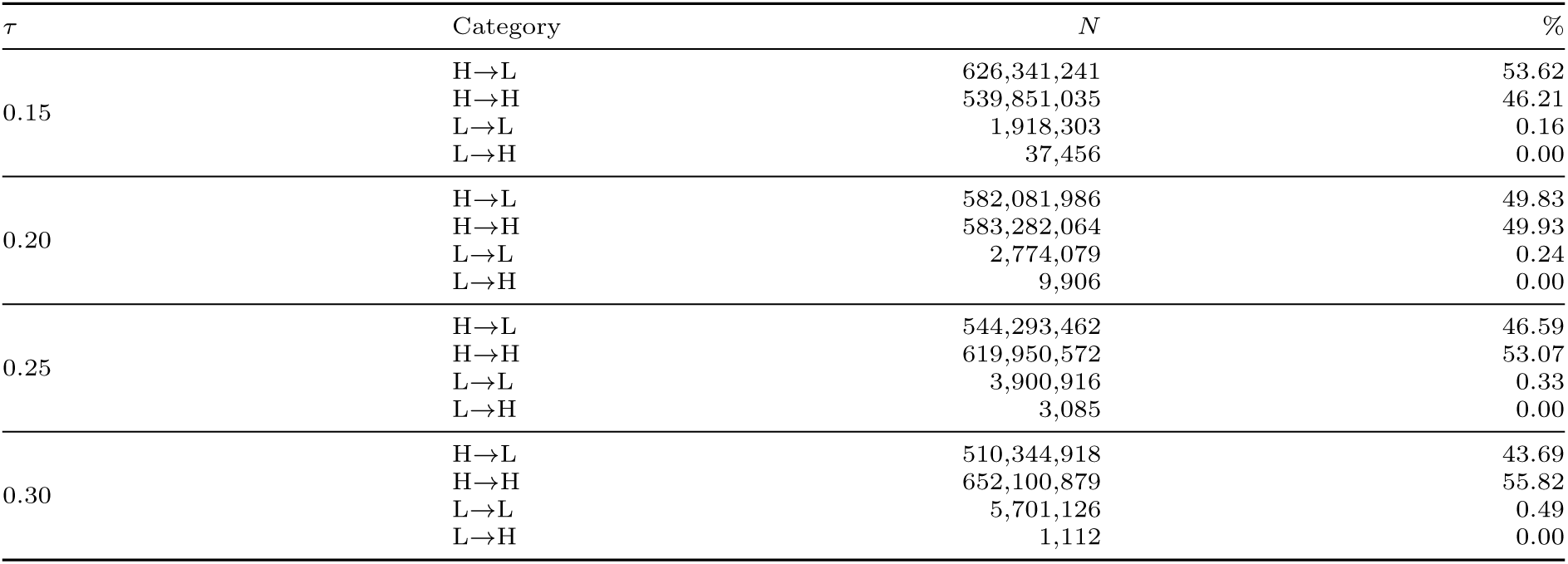
Learnability categorization of Dayhoff-170M-GigaRef training sequences. Sensitivity analysis of threshold parameter *τ* on sequence categorization. Linear fit across 6 checkpoints {1000, 16000, 31000, 46000, 61000, 76000}; reference step 76,000. Global *L*_mean_ = 2.2835 (PPL 9.81). Total sequences: 1,168,148,035 across 574 shards.

**Table S6.**
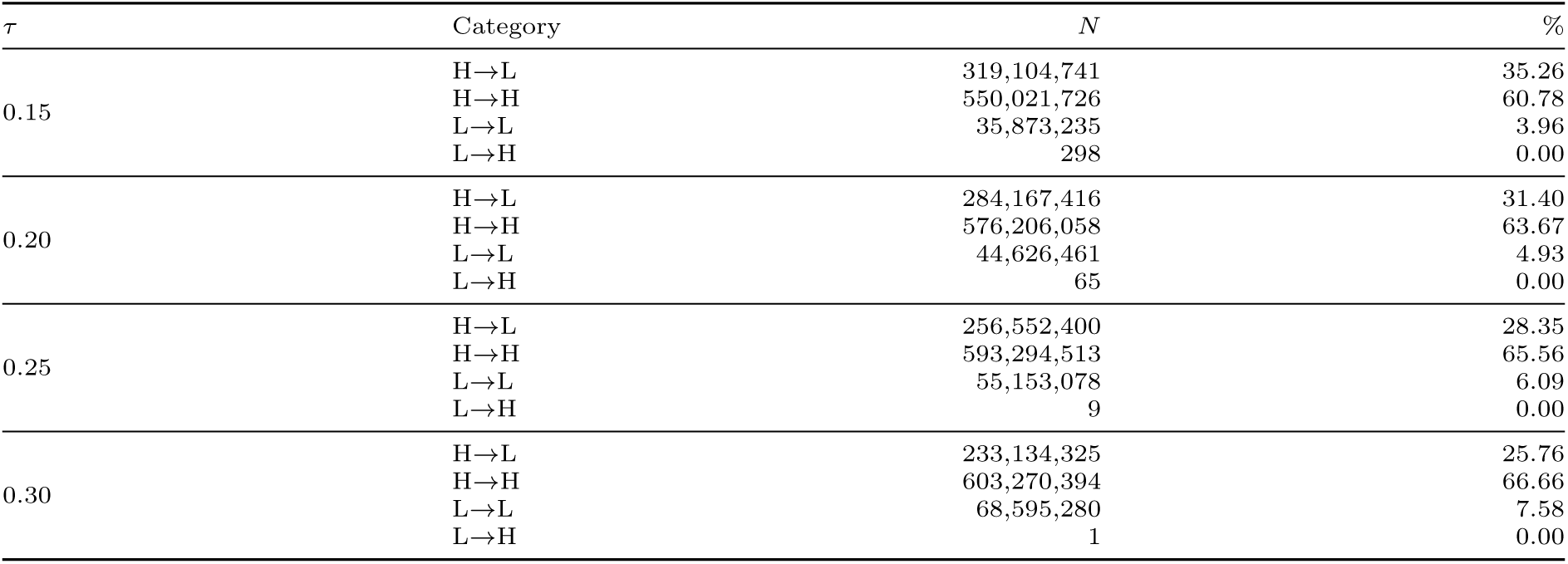
Learnability categorization of Dayhoff-170m-GRsingletons training sequences. Sensitivity analysis of threshold parameter *τ* on sequence categorization. Linear fit across 5 checkpoints {2000, 26000, 50000, 76000, 112000}; reference step 112,000. Global *L*_mean_ = 2.4959 (PPL 12.13). Total sequences: 905,000,000 across 905 shards.

**Table S7.**
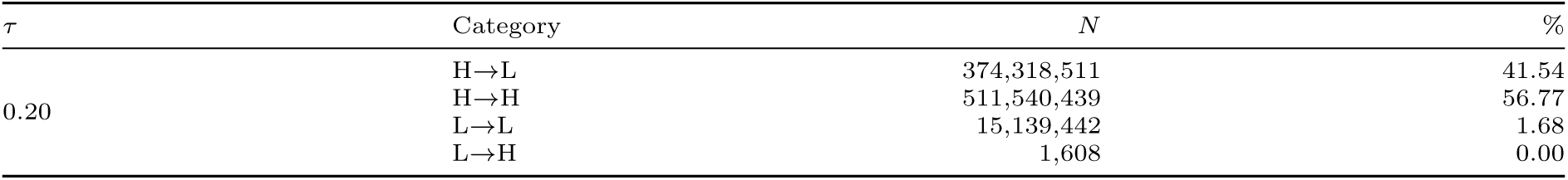
Learnability categorization of Dayhoff-170M-GR evaluated on GigaRef-Singletons. Sequence categorization at threshold parameter *τ* = 0.2. Linear fit across 6 checkpoints {1000, 16000, 31000, 46000, 61000, 76000}; reference step 76,000. Global *L*_mean_ = 2.3682 (PPL 10.68). Total sequences: 901,000,000.

| $\tau$ | Category | $N$ | % |
| --- | --- | --- | --- |
| 0.20 | H→L | 374,318,511 | 41.54 |
|  | H→H | 511,540,439 | 56.77 |
|  | L→L | 15,139,442 | 1.68 |
|  | L→H | 1,608 | 0.00 |

**Table S8.** Learnability categorization of Dayhoff-170m-GRs evaluated on GigaRef-clustered sequences. Sequence categorization at threshold parameter *τ* = 0.2. Linear fit across 5 checkpoints {2000, 26000, 50000, 76000, 112000}; reference step 112,000. Global *L*_mean_ = 2.5204 (PPL 12.43). Total sequences: 1,336,155,089.

| $\tau$ | Category | $N$ | % |
| --- | --- | --- | --- |
| 0.20 | H→L | 412,521,042 | 30.87 |
|  | H→H | 853,189,854 | 63.85 |
|  | L→L | 70,443,411 | 5.27 |
|  | L→H | 782 | 0.00 |

**Table S9.** Cluster size distribution for UR90-based Dayhoff-170M models. Sequence categorization across cluster Cluster sizes for the UR90 and UR90-H-L training corpora. Columns report the number of clusters, total sequences, and the percentage of sequences stratified by learnability category.

| Model | Cluster size | $n_{\text{clusters}}$ | $n_{\text{seqs}}$ | H→H | H→L | L→H | L→L |
| --- | --- | --- | --- | --- | --- | --- | --- |
| Dayhoff-170M-UR90 | 1 | 45,021,254 | 45,021,254 | 61.1% | 34.7% | 0.0% | 4.3% |
|  | 2–10 | 16,456,602 | 55,058,836 | 43.2% | 54.6% | 0.0% | 2.2% |
|  | 11–100 | 2,043,798 | 53,067,115 | 25.6% | 73.2% | 0.0% | 1.2% |
|  | 101–1000 | 138,997 | 28,740,016 | 12.3% | 87.4% | 0.0% | 0.3% |
|  | 1000+ | 1,387 | 2,013,689 | 6.6% | 93.4% | 0.0% | 0.0% |
| Dayhoff-170M-UR90-H-L | 1 | 45,021,254 | 45,021,254 | 81.3% | 16.5% | 0.0% | 2.1% |
|  | 2–10 | 16,456,602 | 55,058,836 | 68.8% | 29.4% | 0.0% | 1.8% |
|  | 11–100 | 2,043,798 | 53,067,115 | 49.4% | 49.0% | 0.0% | 1.6% |
|  | 101–1000 | 138,997 | 28,740,016 | 24.7% | 74.5% | 0.0% | 0.7% |
|  | 1000+ | 1,387 | 2,013,689 | 6.5% | 93.5% | 0.0% | 0.0% |

**Table S10.** Cluster size distribution for GR and GRs Dayhoff-170M models. Sequence categorization across cluster Cluster sizes for the GigaRef and GigaRef-singletons training corpora. Columns report the number of clusters, total sequences, and the percentage of sequences stratified by learnability category. For Dayhoff-170m-GRs, we report performance on training sequences that are singletons (i.e. cluster size 1), and additionally run inference on GigaRef clustered sequences (i.e. cluster size *≥* 1).

| Model | Cluster size | $n_{\text{clusters}}$ | $n_{\text{seqs}}$ | H→H | H→L | L→H | L→L |
| --- | --- | --- | --- | --- | --- | --- | --- |
| Dayhoff-170M-GR | 1 | 60,000,000 | 60,000,000 | 57.6% | 41.6% | 0.0% | 0.9% |
|  | 2–10 | 63,456,065 | 206,146,085 | 50.8% | 48.7% | 0.0% | 0.6% |
|  | 11–100 | 7,668,567 | 222,263,726 | 47.4% | 52.4% | 0.0% | 0.2% |
|  | 101–1000 | 755,989 | 126,540,202 | 52.2% | 47.7% | 0.0% | 0.1% |
|  | 1000+ | 2,390 | 2,974,452 | 59.8% | 40.0% | 0.0% | 0.2% |
| Dayhoff-170m-GRs | 1 | 284,000,000 | 284,000,000 | 62.5% | 31.4% | 0.0% | 6.1% |
|  | 2–10 | 8,572,788 | 27,843,645 | 65.2% | 30.8% | 0.0% | 4.0% |
|  | 11–100 | 1,030,732 | 29,836,884 | 68.8% | 29.7% | 0.0% | 1.5% |
|  | 101–1000 | 102,953 | 17,285,539 | 72.4% | 26.3% | 0.0% | 1.3% |
|  | 1000+ | 364 | 462,882 | 74.8% | 23.2% | 0.0% | 2.0% |

**Table S11.**
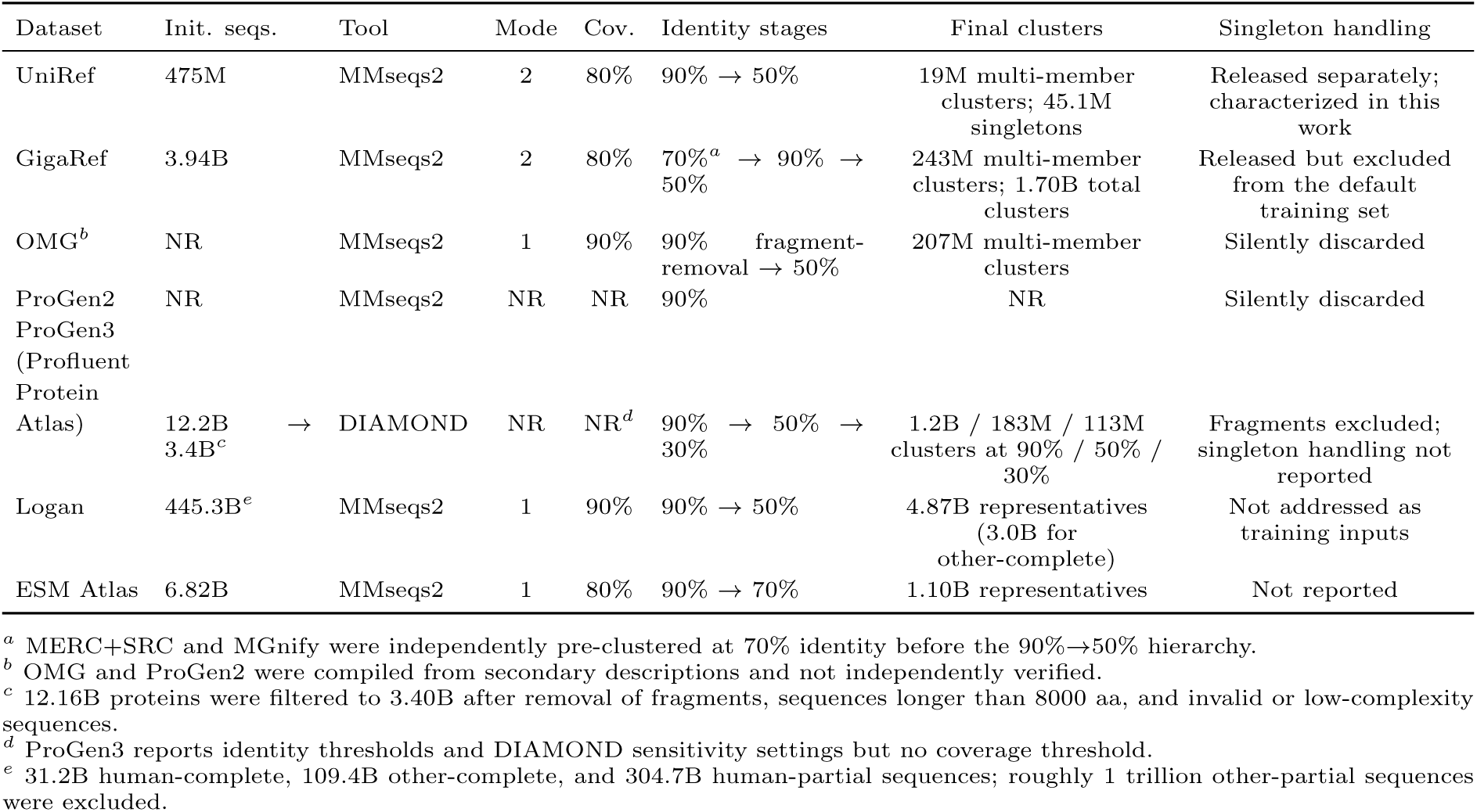
Clustering protocols and singleton handling across large-scale sequence datasets.

## Supplementary Figures

**Figure S1.**
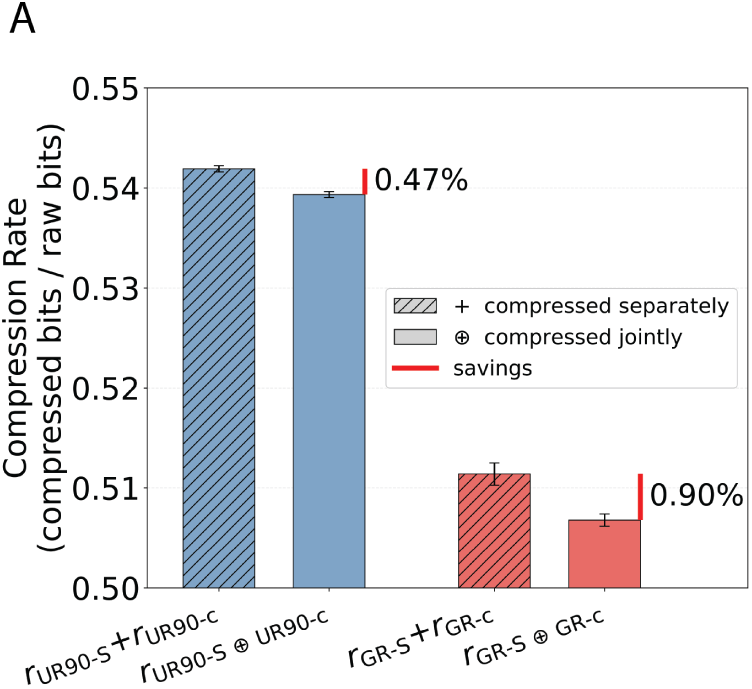
Sequence loss vs. cluster size for multiple PLMs and datasets. **(A)** Compression rate (compressed bits divided by raw bits) for two datasets, UniRef90 (blue) and GigaRef (red). Within each dataset, the hatched bar shows the two subsets compressed separately (+) and the solid bar shows them compressed jointly (⊕). Error bars denote the standard deviation across 50 replicates, and the red mark with its label gives the percentage difference between the two bars.

**Figure S2.**
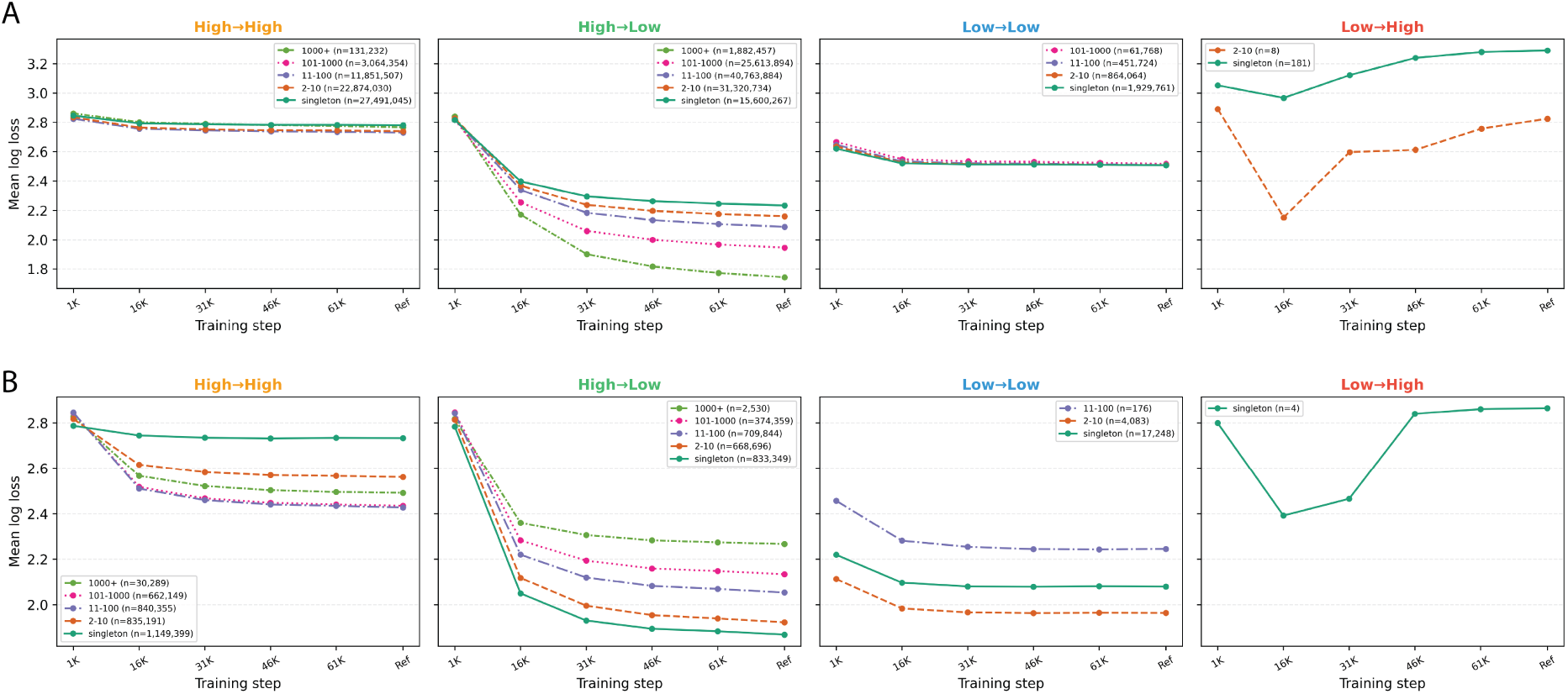
Training loss trajectories of sequences for Dayhoff-170m PLMs stratified by cluster size. Training data for each PLM is (A) UniRef90 (B) GigaRef-clustered. On each panel, *n* denotes the number of sequences in the cluster size bin.

**Figure S3.**
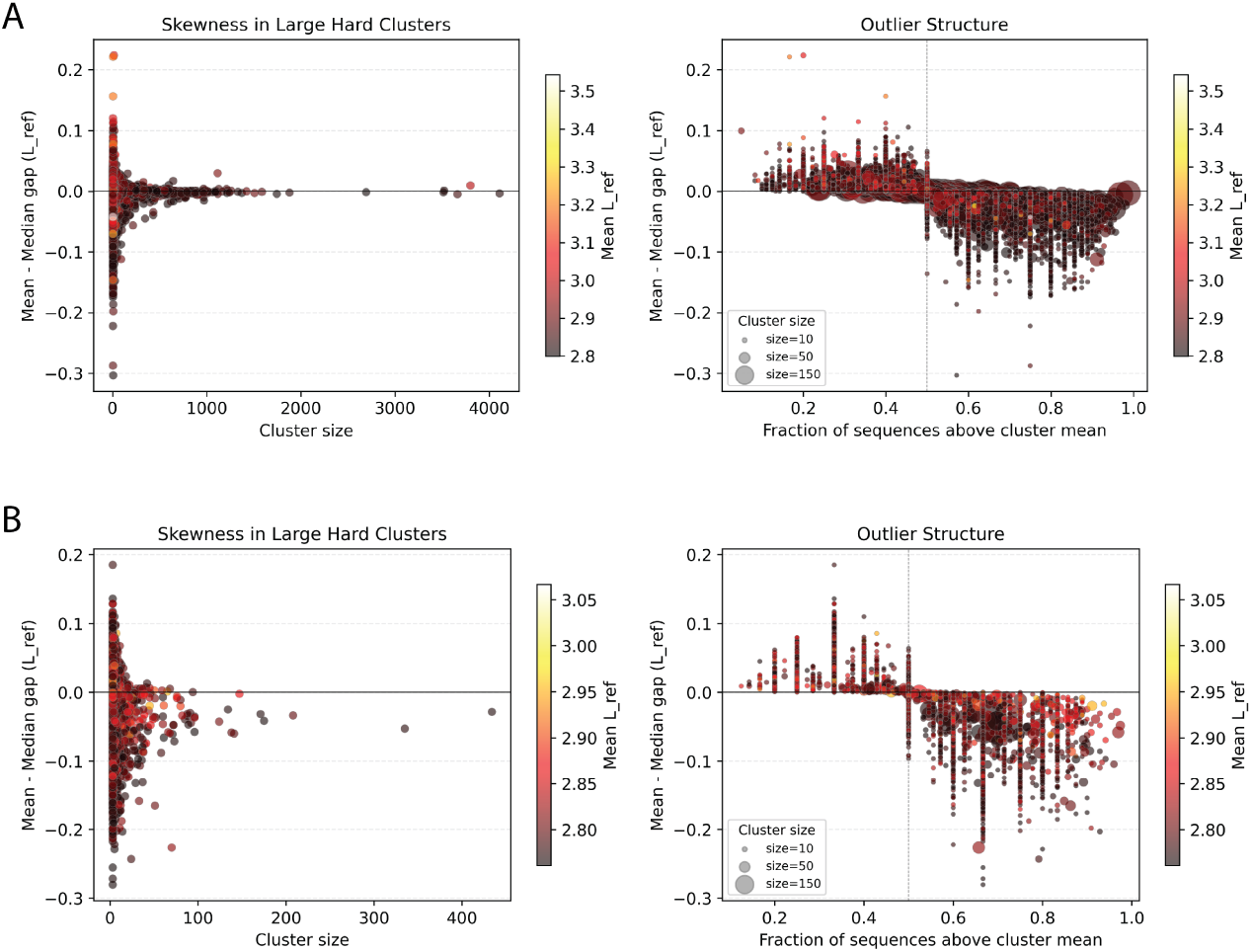
Characterization of large High*→*High-classified clusters by learnability category for (A) Dayhoff-170m-UR90 on UniRef90 sequences (B) Dayhoff-170m-GR on GigaRef-clustered sequences. **Left**: Mean-median gap of final checkpoint sequence loss vs. cluster size, colored by mean final checkpoint sequence loss. **Right**: Mean-median gap of final checkpoint sequence loss vs. fraction of sequences above the cluster mean, with point size scaled by cluster size and color by mean final checkpoint sequence loss.

**Figure S4.**
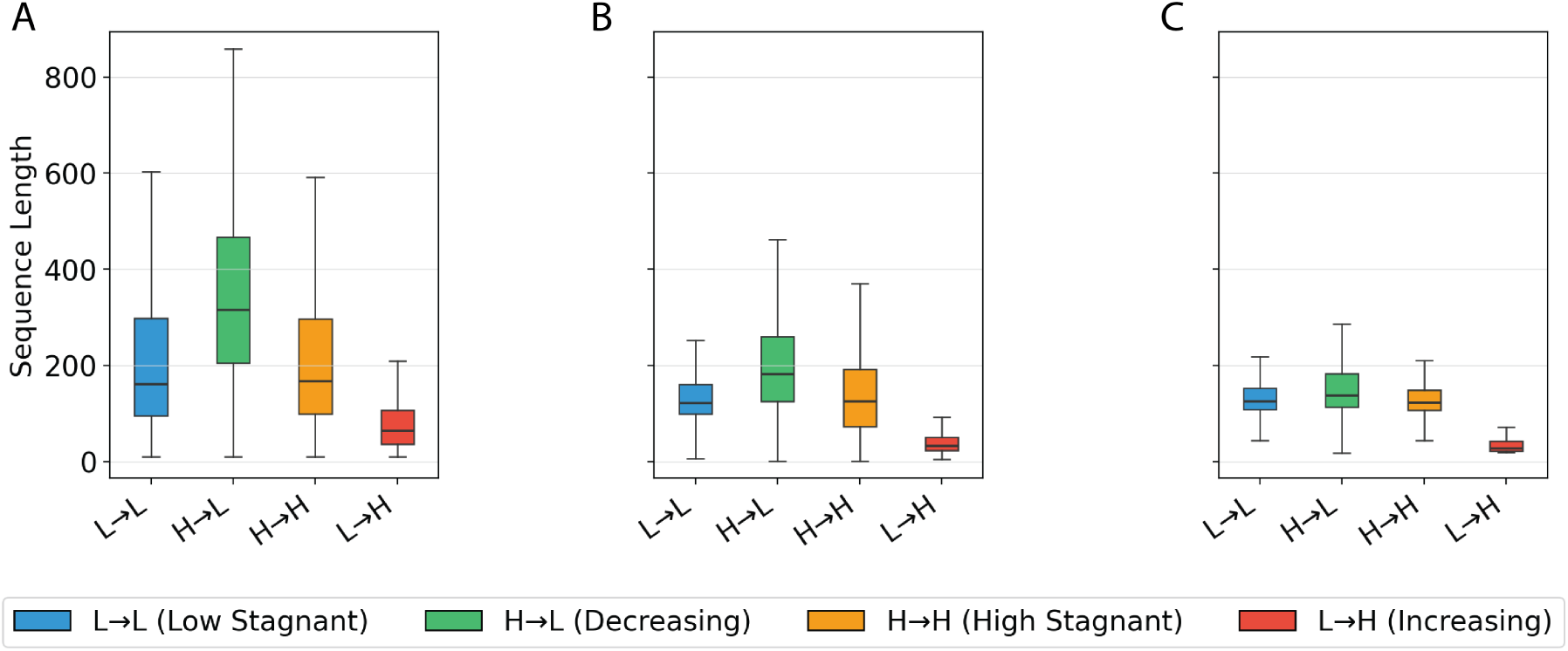
Distribution of sequence lengths of learnability categories for **(A)** Dayhoff-170m-UR90 on UniRef90 sequences **(B)** Dayhoff-170m-GR on GigaRef-clustered sequences **(C)** Dayhoff-170m-GRs on GigaRef-Singleton sequences.

